# CLCluster: a redundancy-reduction contrastive learning-based clustering method of cancer subtype based on multi-omics data

**DOI:** 10.1101/2024.03.07.584010

**Authors:** Hong Wang, Yi Zhang, Wen Li, Zhenlong Wang, Zhen Wei, Mengyuan Yang

## Abstract

Alternative splicing (AS) enables the regulated generation of multiple mRNA and protein products from a single gene. Cancer cells have general, cancer type-specific, and subtype-specific alterations in the splicing process that can have predictive value and contribute to cancer diagnosis, prognosis, and treatment. Currently, multi-omics data have been used to identify the molecular subtype of cancer. However, alternative splicing is rarely used to identify the cancer subtypes. Here, we propose a redundancy-reduction contrastive learning-based method (CLCluster) based on copy number variation, DNA methylation, gene expression, miRNA expression, and alternative splicing for cancer subtype clustering of 33 cancer types. Experimental results demonstrate the superior performance of the proposed CLCluster model in identifying cancer subtypes over the currently available state-of-the-art clustering methods. Moreover, ablation experiments demonstrate the advantages of alternative splicing data for cancer subtyping tasks. We performed multiple analyses for cancer subtype-related AS events, including open reading frame annotation, and RNA binding protein-associated alternative splicing regulation. From our analysis, we identified 2,930 AS events that were associated with patient survival, and ORF analysis showed that 417 of them could cause in-frame and 420 could cause frameshift. we also identified 1,752 RBP-AS regulatory pairs that could be associated with patient survival. Accurate classification of the cancer type using CLCluster, and effective annotation of cancer subtype related AS events can effectively facilitate the identification of patient’s therapeutically targetable AS events.

## 1. Introduction

Cancer stands as a prominent contributor to mortality, posing a substantial obstacle to improving life expectancy globally (1). The intricate and diverse genomic nature of cancer underscores its complexity, with notable variations in tumor responsiveness to the treatment (2). Recognizing distinct cancer subtypes is essential, given their substantial differences in diagnosis, prognosis, and therapeutic approaches (3). With the rapid development of high-throughput molecular technologies, numerous large-scale databases have been created for multi-omics analysis, such as The Cancer Genome Atlas (TCGA) (4). As a genetic disease, cancer develops with abnormalities at multiple levels such as the genome, epigenome, transcriptome, and proteome (5). Utilizing multi-omics data can aid in achieving a systematic and comprehensive understanding of complex biological information at various levels. It supplements the content of all links from genes to phenotypes, contributing to improved treatment and prevention. Additionally, it helps researchers decipher the occurrence and development of cancer at the molecular level (6). Gene expression, miRNA expression, methylation, and copy number variation are commonly used multi-group data examples.

Cancer cells exhibit alterations in the splicing process that are general, specific to certain types of cancer, and distinct to particular subtypes, all of which can have predictive value and contribute to various aspects of cancer progression, including its impact on immune responses. (7). The main alternative splicing patterns are divided into 5 types: Exon skipping, intron retention, mutually exclusive exons, alternative 5′ SS, and alternative 3′ SS. Growing evidence has revealed that misplacing events can change protein function and lead to human diseases. The study of its mechanism will provide important information for the treatment of diseases, including cancer initiation, maintenance, and/or progression (8).

Compared with single-group analysis, multi-group analysis can have a more comprehensive understanding of the mechanism of tumorigenesis. At the present stage, due to complex heterogeneity and high dimensionality, the methods of multi-group data integration and analysis are faced with three fatal problems: (I) heterogeneity in group data; (II) information complementarity among different types and levels of group data; (III) dimensional disaster in group data (9). Many dimensionality reduction approaches have been applied to multi-omics data. SNF(10) fused networks from different data types into a similarity network using a nonlinear strategy. PINS(11) utilized omics data to find strongly connected subgraphs among patient nodes and used similarity-based clustering to analyze merged subgraphs. NEMO(12) clustered the interpatient similarity matrices of multiple omics data to obtain subtyping results. iCluster(13) and NMF(14) model the distribution of each data type and then maximize the likelihood of multi-omics data based on joint latent variables.

However, traditional statistical or mathematical models still face significant challenges in accurately modeling high-dimensional multi-omics data due to their complexity. Deep learning has developed rapidly in recent years and is now widely used in areas such as image recognition, text generation, and language translation (15). In addition to these traditional methods mentioned above, several deep learning-based methods have been developed to solve multi-omics clustering tasks. Chaudhary et. al (16) first applied deep learning to multi-omics data dimensionality reduction using an autoencoder to complete the dimensionality reduction of multi-omics data and then used the k-means algorithm to complete the clustering. Zhao et. al(17) used comparative clustering to implement an end-to-end model that completed both dimensionality reduction and clustering and achieved better clustering results than the autoencoder.

The application of deep learning improves the effectiveness of cancer clustering compared to traditional methods, and comparative learning in deep learning is best able to obtain a meaningful low-dimensional representation of the data. Contrastive learning is a form of self-supervised learning, a successful approach to SSL is to learn embeddings that are invariant to distortions of the input sample. Although contrastive learning can achieve better dimensionality reduction than other methods, it has problems with the existence of trivial constant solutions and model collapse (18). To address the above problem we introduce CLCluster, which is a model that combines comparative learning and mean-shift cluster. CLCluster is a unique model that differs from other contrast learning models. Unlike them, it doesn’t use negative examples. Instead, it applies redundancy-reduction, a principle that was first proposed in neuroscience, to self-supervised learning. This approach helps to avoid getting trivial constant solutions and reduces the probability of model collapse. Mean-shift enables automatic clustering, avoiding the need to preset the number of clustering categories like k-means. This reduces the impact of human intervention on clustering results. To evaluate the performance of CLcluster, we compared its performance with that of 7 state-of-the-art multi-omics data clustering methods on 33 datasets from TCGA. The results show that CLcluster achieves competitive results on most datasets. On this basis, we used CLcluster to investigate the effect of introducing alternative splicing data on the clustering effect of cancer subtypes. In addition, we performed a series of biological analyses to demonstrate the biological significance of the subtypes obtained by CLcluster, as well as screening for potential biomarkers associated with alternative splicing.

## 2. Materials and Methods

### 2.1 Data

We first obtained 33 cancers’ multi-omics data from the TCGA portal (https://tcga-data.nci.nih.gov/tcga/) (4). These data sets include gene expression, DNA methylation, miRNA expression, copy number alterations (CNA), and patient clinical information. Then, we collected alternative splicing events information on TCGA from the study by Kahles et al. (https://bioinformatics.mdanderson.org/TCGASpliceSeq/index.jsp) (19). Our data preprocessing methodology comprises three principal steps, influenced by insights gleaned from earlier research(10). For DNA methylation, we mapped the methylated site of each gene and computed the average methylation beta value for each gene. The biological features including gene expression, miRNA expression, DNA methylation, miRNA expression, and alternative splicing were filtered if they had null values in more than 20% of patients. The samples were removed if missing across more than 20% of features. Then, we used the impute function from the R impute package(20) to fill out the missing values. The biology feature numbers used to train the CLCluster of each cancer are shown in **Table 1**.

**Table 1.**
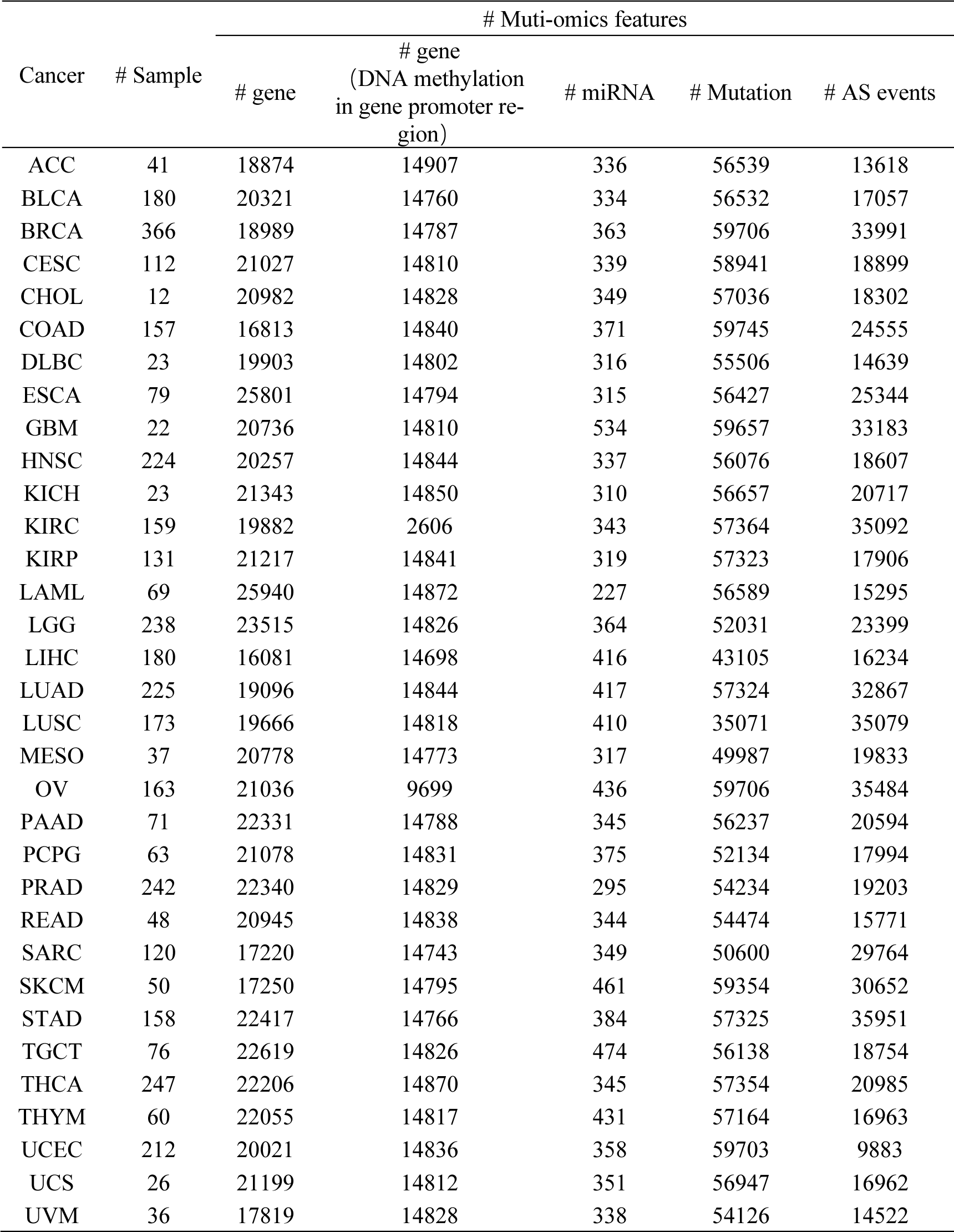
Data and sample size of CLCluster.

| Cancer | # Sample | # Multi-omics features |  |  |  |  |
| --- | --- | --- | --- | --- | --- | --- |
|  |  | # gene | # gene<br>(DNA methylation<br>in gene promoter re-<br>gion) | # miRNA | # Mutation | # AS events |
| ACC | 41 | 18874 | 14907 | 336 | 56539 | 13618 |
| BLCA | 180 | 20321 | 14760 | 334 | 56532 | 17057 |
| BRCA | 366 | 18989 | 14787 | 363 | 59706 | 33991 |
| CESC | 112 | 21027 | 14810 | 339 | 58941 | 18899 |
| CHOL | 12 | 20982 | 14828 | 349 | 57036 | 18302 |
| COAD | 157 | 16813 | 14840 | 371 | 59745 | 24555 |
| DLBC | 23 | 19903 | 14802 | 316 | 55506 | 14639 |
| ESCA | 79 | 25801 | 14794 | 315 | 56427 | 25344 |
| GBM | 22 | 20736 | 14810 | 534 | 59657 | 33183 |
| HNSC | 224 | 20257 | 14844 | 337 | 56076 | 18607 |
| KICH | 23 | 21343 | 14850 | 310 | 56657 | 20717 |
| KIRC | 159 | 19882 | 2606 | 343 | 57364 | 35092 |
| KIRP | 131 | 21217 | 14841 | 319 | 57323 | 17906 |
| LAML | 69 | 25940 | 14872 | 227 | 56589 | 15295 |
| LGG | 238 | 23515 | 14826 | 364 | 52031 | 23399 |
| LIHC | 180 | 16081 | 14698 | 416 | 43105 | 16234 |
| LUAD | 225 | 19096 | 14844 | 417 | 57324 | 32867 |
| LUSC | 173 | 19666 | 14818 | 410 | 35071 | 35079 |
| MESO | 37 | 20778 | 14773 | 317 | 49987 | 19833 |
| OV | 163 | 21036 | 9699 | 436 | 59706 | 35484 |
| PAAD | 71 | 22331 | 14788 | 345 | 56237 | 20594 |
| PCPG | 63 | 21078 | 14831 | 375 | 52134 | 17994 |
| PRAD | 242 | 22340 | 14829 | 295 | 54234 | 19203 |
| READ | 48 | 20945 | 14838 | 344 | 54474 | 15771 |
| SARC | 120 | 17220 | 14743 | 349 | 50600 | 29764 |
| SKCM | 50 | 17250 | 14795 | 461 | 59354 | 30652 |
| STAD | 158 | 22417 | 14766 | 384 | 57325 | 35951 |
| TGCT | 76 | 22619 | 14826 | 474 | 56138 | 18754 |
| THCA | 247 | 22206 | 14870 | 345 | 57354 | 20985 |
| THYM | 60 | 22055 | 14817 | 431 | 57164 | 16963 |
| UCEC | 212 | 20021 | 14836 | 358 | 59703 | 9883 |
| UCS | 26 | 21199 | 14812 | 351 | 56947 | 16962 |
| UVM | 36 | 17819 | 14828 | 338 | 54126 | 14522 |

### 2.2 Construction of CLCluster model

#### 2.2.1 Construction of redundancy-reduction contrastive learning model

CLCluster (shown in **Figure 1**) is a redundancy-reduction contrastive learning clustering method based on multi-omics datasets for cancer subtyping. The method performs feature extraction by redundancy-reduction contrast learning model. For the extracted features, after introducing survival information for further dimensionality reduction, clustering is performed using Mean-Shift to obtain cancer subtypes. To prevent model collapse and not require negative examples or asymmetric structures, we employ a unique loss function for Redundancy-reduction contrastive learning.

**Figure 1.**
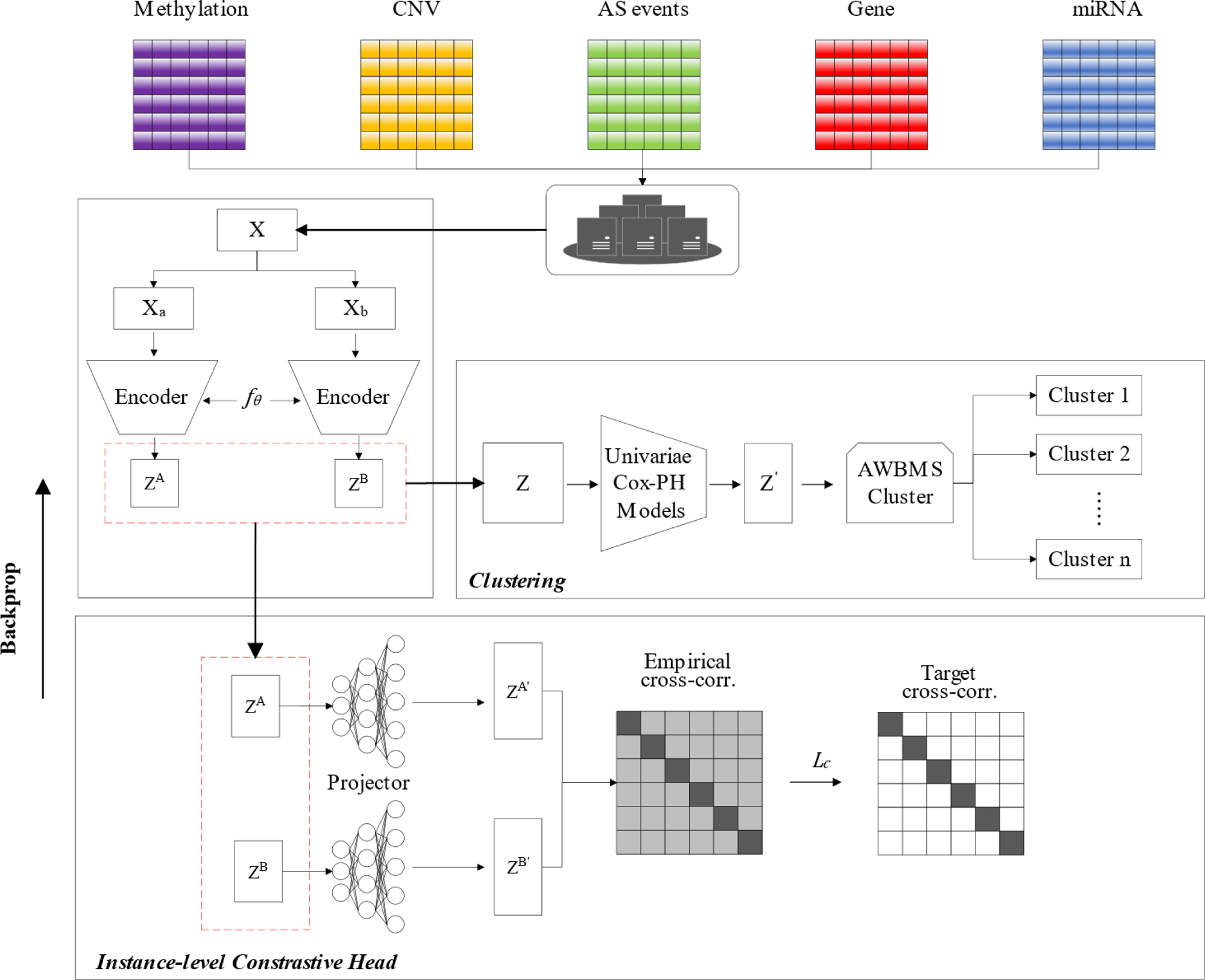
The framework of CLCluster. First, the five multi-omics datasets were merged to form a single matrix input. The data pairs are augmented using data augmentation and embeddings are extracted from the data expansion using a shared autoencoder. To enhance the learning process, a shared neural network is utilized to project the embeddings. Instance-level comparative learning is then carried out based on redundancy-reduction loss. Following the training phase, the encoder part of the autoencoder undergoes survival analysis to eliminate features that are not related to survival. Next, these features are automatically clustered using mean-shift to obtain different cancer subtypes. Contrastive Learning via Redundancy-Reduction.

CLCluster operates on a joint embedding of augmentation data. Specifically, given a data instance *x*, two stochastic data transformations *T^a^, T^b^* sampled from the same family of augmentations *T* are applied to it, resulting in two correlated samples denoted as 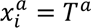 and 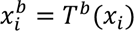. A proper choice of augmentation strategy is essential to achieve a good performance in downstream tasks. For this research, a method of improving data quality is implemented, which involves adding noise to the existing data. Specifically, to enhance the data, we add standard Gaussian noise to the original matrix. One shared deep neural network f(·) is used to extract features from the augmented sample 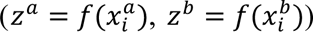. Specifically, our method adopts a four-layer Variational AutoEncoder(VAE). The hidden layer of VAE will be used as a low-dimensional embedding obtained by dimensionality reduction for downstream clustering of cancer subtypes. We avoid directly conducting contrastive learning on the feature matrix to enhance contrastive learning performance. Instead, we stack a three-layer nonlinear MLP g(·) to map the feature matrix to a subspace via 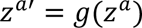 where the redundancy-reduction contrastive loss is applied. To effectively solve the model collapse problem, the CLCluster loss function does not use negative examples, but redundancy-reduction loss functions *L_CL_*:

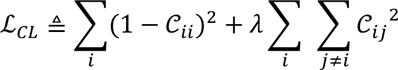

λ is a positive constant that balances the importance of the first and second terms in the loss function. *C* is the cross-correlation matrix computed between the outputs of the two identical networks along the batch dimension:

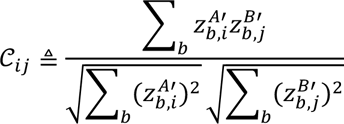

where *b* indexes batch samples and *i*, *j* index the vector dimension of the networks’ outputs. *C* is a square matrix with size the dimensionality of the network’s output, and with values comprised between −1 (perfect anti-correlation) and 1 (perfect correlation).

The invariance term of the objective, by trying to equate the diagonal elements of the cross-correlation matrix to 1, makes the embedding invariant to the distortions applied. The redundancy-reduction term, by trying to equate the off-diagonal elements of the cross-correlation matrix to 0, decorates the different vector components of the embedding. This decorrelation reduces the redundancy between output units so that the output units contain non-redundant information about the sample. Compared to INFONCE-based methods, our method is not restricted by negative examples, can achieve good results in small batches of inputs, and is friendly to high-dimensional input data.

#### 2.2.2 Transformed feature selection and automatic clustering based on Mean Shift Algorithm

Contrastive Learning via redundancy reduction reduced the initial number of features to 500 new features obtained from the bottleneck layer of VAE. Next, for each of these transformed features produced by the autoencoder, we built a univariate Cox proportional hazards (Cox-PH) model and selected features from which a significant Cox-PH model is obtained (log-rank P < 0.05). We then used these reduced new features to cluster the samples using the Mean Shift Algorithm (MS). Most of the algorithms such as k-means and its variants (21), hierarchical clustering(Carlsson and Memoli 2010), spectral clustering(23), and density-based methods(24), inherently use the number of clusters (*k*) as an input. However, for real-world data, k may not be known beforehand. Determining k from the dataset itself has long been an open problem. To find the number of clusters automatically and to learn various properties of the feature space, we have resorted to the mean shift (MS) paradigm (25).

### 2.3 Parameter settings of CLcluster

The model is developed with Python 3.10, Pytorch 2.0.1, and CUDA11.7. Since the dataset structure of 33 cancers is similar, we chose the dataset of hepatocellular carcinoma in the training of CLCluster. Compared with other cancer data sets, the data amount and feature number of hepatocellular carcinoma data sets are moderate and representative. Because contrastive learning is a self-supervised deep learning model, there is no explicit label to evaluate the performance of the model, so we divide the training effect into two stages to verify. In the first stage, we output the loss value of each training, draw the loss image, and record the parameters that the loss value gradually decreases and the overall trend of the image is smooth. In the second stage, we use the features obtained from the trained model to extract and cluster the survival features mentioned above. For the clustering results, we first conduct a single-factor survival analysis of the clustering results combined with the survival information to see if there is a survival difference between the clustering results. at the same time, we also use the t-SNE (t-Distributed Stochastic Neighbor Embedding) algorithm to visualize the clustering results to see the clustering distribution of the samples in the space. Finally, the model training parameters with a high degree of dispersion between subtypes and survival related to the subtypes were recorded. After completing the training on hepatocellular carcinoma, we apply the model to the remaining 32 cancer data sets, and fine-tune the model according to the specific conditions of each data set, and finally get a training parameter that can achieve better results on all data sets. To optimize the model performance, we have fine-tuned the following parameters: tuning batch size to optimize Contrastive Learning performance; altering feature dimension to change the size of the feature space for keeping data information; adjusting the learning rate to optimize the speed of model converges and changing training epoch to set the appropriate training time for the model. The parameter selection is shown in **Table 2**. For the clustering task, we adjust the parameters bandwidth for each data set, our parameter adjustment range is 0.98-0.05, and the step size is 0.01. For the interval of the result oscillation, we further adjust the step size to 0.01, traverse all the values in the interval, and finally get the optimal value, as shown in **Table 3**.

**Table 2.** Hyperparameter of CLCluter.

| Hyperparameter | Selection range | Optimal value |
| --- | --- | --- |
| batch_size | 16,32,64,128 | 16 |
| feature dimen-<br>tion | 50,100,150,200 | 50 |
| learning rate | 0.001,0.0001 | 0.0001 |
| epochs | [100,1000],<br>step=100 | 600 |

**Table 3.** Parameter of meanshift.

| Cancer | Bandwidth | Cancer | Bandwidth |
| --- | --- | --- | --- |
| KIRC | 0.2 | CHOL | 0.255 |
| LIHC | 0.2 | DLBC | 0.26 |
| SARC | 0.2 | KICH | 0.25 |
| LUSC | 0.25 | KIRP | 0.3 |
| COAD | 0.32 | MESO | 0.185 |
| OV | 0.22 | PAAD | 0.32 |
| STAD | 0.31 | READ | 0.575 |
| GBM | 0.37 | UCS | 0.57 |
| SKCM | 0.33 | LUAD | 0.24 |
| HNSC | 0.29 | BRCA | 0.28 |
| LGG | 0.27 | ESCA | 0.285 |
| UCEC | 0.21 | TGCT | 0.32 |
| THCA | 0.29 | THYM | 0.35 |
| PRAD | 0.43 | UVM | 0.23 |
| ACC | 0.24 | LAML | 0.255 |
| BLCA | 0.4 | PCPG | 0.315 |
| CESC | 0.27 |  |  |

### 2.4 Performance metrics

Four metrics were used to evaluate the performance of different algorithms. (i) *Silhouette score:* the silhouette value measures how similar an object is to its cluster (cohesion) compared to other clusters (separation). The silhouette ranges from −1 to +1, where a high value indicates that the object is well-matched to its cluster and poorly matched to neighboring clusters. If most objects have a high value, then the clustering configuration is appropriate. If many points have a low or negative value, then the clustering configuration may have too many or too few clusters (26). (ii) *Concordance index(C-index):* the concordance index (C-index) can be seen as the fraction of all pairs of individuals whose predicted survival times are correctly ordered(27) and is based on Harrell C statistics(28). The C-index value greater than 0.7 indicates a good prediction result of the model, while a value less than 0.5 indicates high randomness. (iii) Davies *Bouldin index (DBI)*: Davies Bouldin score is a metric for evaluating clustering algorithms. (29) The Davies Bouldin score calculates the sum of the average distance of any two categories divided by the center distance of two clusters to obtain the maximum value. Lower index values indicate a better clustering result. The index is improved (lowered) by increased separation between clusters and decreased variation within clusters. (iv) *Log-rank P value of COX Proportional Hazards (COX-PH):* Taking survival outcome and survival time as dependent variables, COX-PH(30) can analyze the influence of many factors on survival time at the same time.

### 2.5 Analysis of subtype prognosis-related alternative splicing events

First, we performed ANOVA analysis to identify the subtype-related AS events with a P-value less than 0.05 for different subtypes of AS. Then the AS events with the difference in PSI between the most favorable and least favorable prognostic subtypes was more than 0.1 and the p-value smaller than 0.05 were selected as subtype related AS events. Next, Pearson correlation analysis was performed between the AS events and subtype prognosis stag, then the correlation of Pearson was greater than 0.3 and p-value <0.05 were identified as subtype prognosis-related alternative splicing events.

### 2.6 Open reading frame variation annotation of alternative splicing events

For each AS event, we assigned each event to specific regions of the transcript, namely the coding sequence (CDS), 5’ untranslated region (5’UTR), and 3’ untranslated region (3’UTR). Then, for AS events in CDS, we examined isoform sequences’ open reading frames. We reported transcripts as ‘in-frame’ if the number of sequences was a multiple of three after removing the skipped exon sequence. One or two nucleotide insertions were classified as ‘frame-shift’.

### 2.7 Analysis of subtype-related RNA binding protein and RNA binding protein-associated AS events

For ∼1500 RBP(32), we performed ANOVA analysis and identified the subtype-related RNA binding proteins(RBPs). Then, Pearson analysis was performed between the expression of RBP and the PSI value of AS events to identify the RBP-associated AS events for each cancer type (|ρ| >0.3 and p-value<0.05).

## 3 Results

### 3.1 The performance of CLCluster compared to other models

We conduct the clustering performance comparison of CLCluster against 7 state-of-the-art methods for multi-omics data integration including NEMO(12), Subtype-DCC(17), AE(16), SNF(10), iCluster(33), NMF(14), and PINS(11). We evaluate the clustering results of the 7 methods using the multi-group dataset from CLCluster and the previously mentioned evaluation criteria. Since Subtype-DCC and AE use deep learning to reduce the dimension of the original data, C-index is only used to evaluate the results of Subtype-DCC and AE. The results are shown in **Table S1**.

Silhouette and Davies Bouldin scores can characterize the differences between subtypes. CLCluster showed superior silhouette and DBI scores in most cancers compared to other methods (**Figure 2A, B**). There are distinct variations among the subtypes acquired through clustering. Compared with other methods, CLCluster, Subtype-DCC, and AE have higher silhouette and DBI scores. This shows that the difference between subtypes obtained by reducing dimension and clustering by deep learning is stronger. This demonstrates a stronger difference between subtypes obtained through deep learning.

**Figure 2.**
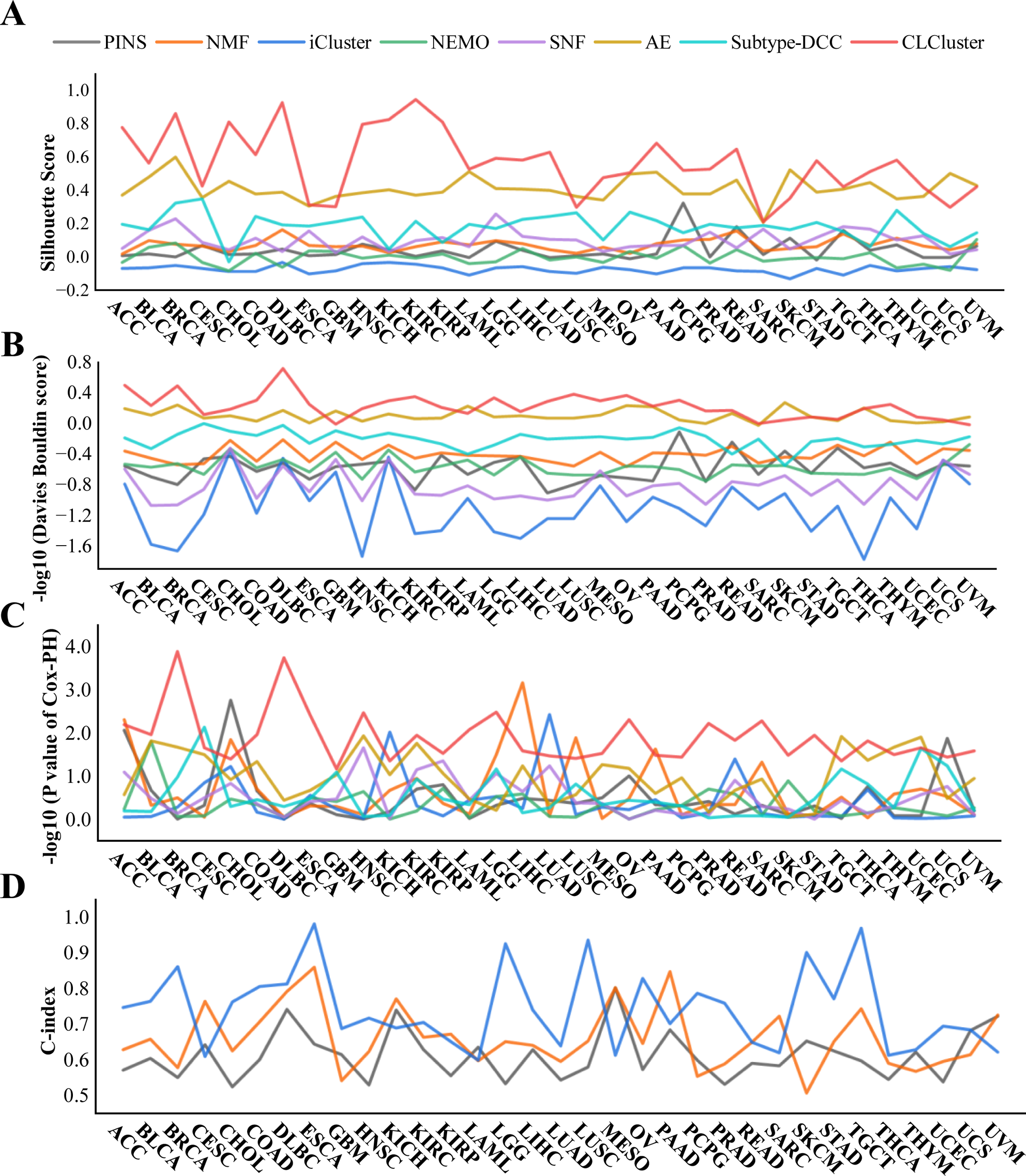
Performance indexes between CLCluster and seven models on 33 cancer data sets. **(A)** Silhouette Score. **(B)** −log10 (Davies Bouldin score). **(C)** −log10 (P value of Cox-PH regression). **(D)** C-index.

The p-value of Cox-PH regression is the result of Cox-PH analysis characterized by clustered subtypes. This value indicates whether there is a correlation between the clustered subtypes and the survival time of the patients (p<0.05 proves a correlation). To facilitate the observation, we have taken the Log-rank of the P value and added a negative sign, at this time the larger the −Log-rank P value, the stronger the correlation is proved to be. As can be seen from **Figure 2C**, there is a correlation between the subtypes obtained by CLCluster in all cancers and patient survival, and the scores of the subtypes obtained by CLCluster in most cancers are higher than those of other methods. This suggests that the subtypes obtained from CLCluster clustering are instructive for patient prognosis.

We performed COX-PH analysis on the features obtained from CLCluster dimensionality reduction to obtain the C-index value. This value indicates the strength of the correlation between the features obtained from dimension reduction and the patient’s survival time. A higher value indicates a stronger correlation. As can be seen from **Figure 2D and Table S1**, the features obtained from CLCluster’s dimensionality reduction for all cancers are consistent with patient survival (C-index value > 0.5), 19 results being highly consistent (C-index value > 0.7). This suggests that CLCluster can extract features that have significance for patient survival and that the extracted features can guide prognosis to some extent.

### 3.2 The performance of CLCluster is enhanced by the introduction of AS data

To demonstrate the advantages of alternative splicing data for cancer subtyping tasks, we performed ablation experiments about alternative splicing data on 33 cancer datasets. We removed the alternative splicing data from each cancer dataset while keeping the sample size constant. Then, we clustered each cancer using CLCluster and analyzed the clustering results using the metrics described above. The results showed that after the introduction of AS data, more than one index of all cancers was improved, and more than 31 cancers were better than those without AS data in more than half of the evaluation results (**Figure 3A-D**). From **Figure 3**, it can be seen that all four metrics (Silhouette score, C index, DBI index, −LOG10(p-value)) of OV (“with AS” vs “without AS”: Silhouette score 0.616 vs 0.447, C-index 0.806 vs 0.622, −LOG10(p-value) 1.957 vs 1.519, −LOG10(DBI) 0.34 vs 0.044), KIRC (“with AS” vs “without AS”: Silhouette score 0.781 vs 0.244, C-index 0.784 vs 0.642, −LOG10(p-value) 2.189 vs 1.438, −LOG10(DBI) 0.5 vs −0.066), BRCA (“with AS” vs “without AS”: Silhouette score 0.578 vs 0.394, C-index 0.773 vs 0.552, −LOG10(p-value) 1.953 vs 1.321, −LOG10(DBI) 0.09 vs −0.048), SARC (“with AS” vs “without AS”: Silhouette score 0.862 vs 0.403, C-index 0.862 vs 0.668, −LOG10(p-value) 3.898 vs 1.832, −LOG10(DBI) 0.385 vs 0.183), ACC (“with AS” vs “without AS”: Silhouette score 0.594 vs 0.268, C-index 0.927 vs 0.68, −LOG10(p-value) 2.481 vs 2.012, −LOG10(DBI) 0.335 vs 0.01), PAAD (“with AS” vs “without AS”: Silhouette score 0.53 vs 0.325, C-index 0.76 vs 0.741, −LOG10(p-value) 2.224 vs 1.421, −LOG10(DBI) 0.163 vs −0.097), and THYM (“with AS” vs “without AS”: Silhouette score 0.582 vs 0.371, C-index 0.629 vs 0.519, −LOG10(p-value) 1.896 vs 1.549, −LOG10(DBI) 0.252 vs 0.053) are significantly improved after the introduction of AS data, and the clustering effect is significantly improved. After the introduction of AS data, all the evaluation indexes for these seven cancers increased by more than 30%. The DBI index of OV increased by a factor of 7, and all indexes of SARC increased by a factor of 2. Overall, the assessment scores of most cancers decreased after excluding AS data. The result proved the importance of alternative splicing and guided our subsequent screening of biomarkers (**Figure 3, Table S2**).

**Figure 3.**
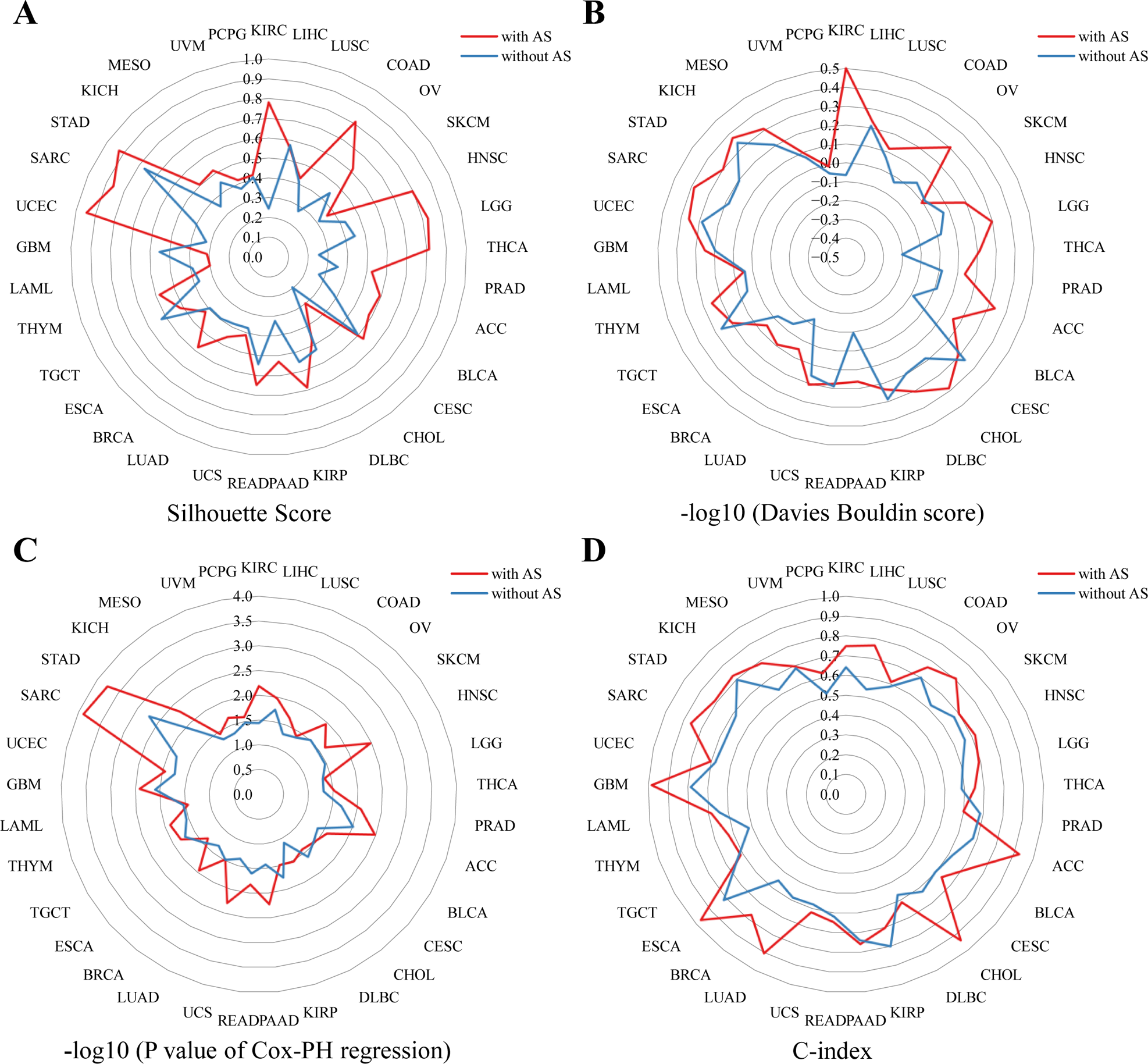
The effect of introducing AS data on model performance. **(A)** −log10 (P value of Cox-PH regression). **(B)** Silhouette score. **(C)** C-index score. **(D)** −log10 (Davies Bouldin score).

### 3.3 CLCluster analysis has identified distinct subtypes that are correlated with cancer prognosis

Patients with different cancer subtypes often have varying prognoses, which can guide clinical treatment decisions. We utilized CLCluster to dimension reduction and cluster multi-group data from 33 cancers, identifying subtypes for each cancer. The results of subtype clustering are shown in **Table 4, Table S3**. To verify the effects of the prognosis predictions of different cancer subtypes, we plotted survival curves of the CLCluster on 33 cancer datasets (**Figure S1**). According to **Figure S1**, the cancer subtypes identified by our method on the other 33 datasets show significantly different survival curves. A significant difference in survival curves was observed among most cancer subtypes, like LIHC, READ, and BLCA. This difference increased over time, indicating varying survival probabilities. However, the clustering effect of CLCluster on some cancers with a small number of samples needs to be improved. The survival curve cannot differentiate patient survival status or reflect prognosis, such as CHOL, and GBM. Based on the embedding learned by our model, we reduced the dimensionality of hidden layer factors. We used t-SNE(34) to visualize corresponding clusters to study the subtypes identified by the CLCluster. According to **Figure S2**, BRCA was divided into 4 subtypes, LIHC was divided into 4 subtypes, and LUAD was divided into 4 subtypes, proving that the model learned a meaningful latent representation. Overall, our findings suggest that CLCluster can effectively identify cancer subtypes associated with prognosis and demonstrate distinct survival patterns across various cancer types.

**Table 4.** The patient number for each Subtype of 32 cancers.

| Cancer | Number of Subtypes | Patient number in each Subtype |
| --- | --- | --- |
| ACC | 5 | ACC-1(20); ACC-2(8); ACC-3(5); ACC-4(5); ACC-5(2) |
| BLCA | 2 | BLCA-1(108); BLCA-2(68) |
| BRCA | 4 | BRCA-1(213); BRCA-2(76); BRCA-3(44); BRCA-4(19) |
| CESC | 4 | CESC-1(35); CESC-2(35); CESC-3(25); CESC-4(17) |
| CHOL | 4 | CHOL-1(5); CHOL-2(1); CHOL-3(1); CHOL-4(1) |
| COAD | 3 | COAD-1(58); COAD-2(56); COAD-3(38) |
| DLBC | 3 | DLBC-1(8); DLBC-2(7); DLBC-3(1) |
| ESCA | 2 | ESCA-1(43); ESCA-2(21) |
| GBM | 4 | GBM-1(11); GBM-2(3); GBM-3(1); GBM-4(1) |
| HNSC | 4 | HNSC-1(155); HNSC-2(26); HNSC-3(23); HNSC-4(20) |
| KICH | 5 | KICH-1(7); KICH-2(4); KICH-3(2); KICH-4(2); KICH-5(1) |
| KIRC | 4 | KIRC-1(90); KIRC-2(35); KIRC-3(13); KIRC-4(6) |
| KIRP | 3 | KIRP-1(73); KIRP-2(38); KIRP-3(17) |
| LAML | 2 | LAML-1(51); LAML-2(13) |
| LGG | 4 | LGG-1(104); LGG-2(74); LGG-3(28); LGG-4(26) |
| LIHC | 4 | LIHC-1(108); LIHC-2(43); LIHC-3(16); LIHC-4(9) |
| LUAD | 4 | LUAD-1(119); LUAD-2(74); LUAD-3(29); LUAD-4(2) |
| LUSC | 4 | LUSC-1(54); LUSC-2(44); LUSC-3(35); LUSC-4(35) |
| MESO | 4 | MESO-1(17); MESO-2(7); MESO-3(5); MESO-4(3) |
| OV | 4 | OV-1(54); OV-2(49); OV-3(32); OV-4(25) |
| PAAD | 3 | PAAD-1(41); PAAD-2(16); PAAD-3(7) |
| PCPG | 2 | PCPG-1(31); PCPG-2(25) |
| PRAD | 3 | PRAD-1(114); PRAD-2(69); PRAD-3(57) |
| READ | 2 | READ-1(40); READ-2(8) |
| SARC | 5 | SARC-1(70); SARC-2(24); SARC-3(17); SARC-4(8); SARC-5(1) |
| SKCM | 4 | SKCM-1(20); SKCM-2(16); SKCM-3(11); SKCM-4(1) |
| STAD | 4 | STAD-1(132); STAD-2(18); STAD-3(1); STAD-4(1) |
| TGCT | 4 | TGCT-1(22); TGCT-2(17); TGCT-3(16); TGCT-4(15) |
| THCA | 3 | THCA-1(140); THCA-2(63); THCA-3(37) |
| THYM | 2 | THYM-1(49); THYM-2(7) |
| UCEC | 4 | UCEC-1(118); UCEC-2(51); UCEC-3(24); UCEC-4(15) |
| UCS | 5 | UCS-1(12); UCS-2(6); UCS-3(3); UCS-4(2); UCS-5(1) |
| UVM | 4 | UVM-1(14); UVM-2(8); UVM-3(8); UVM-4(2) |

### 3.4 Subtype-related AS events provided potential therapeutic targets

Substantial preclinical work has identified a variety of small-molecule compounds as well as genetic and other approaches to target the spliceosome or its products with potential therapeutic effects (7). Since exon skip (ES) results in the loss of functional domains/sites or frameshifting of the open reading frame (ORF), skipped exons have been recognized as therapeutic targets (35, 35, 36). The ES event causes ORF translocation, which in turn leads to protein variation (37). Based on the cancer subtypes obtained by CLCluster, we performed ANOVA analysis and identified 41831 AS events that showed different PSI values across cancer subtypes of 30 cancers. Person association study found 2,921 AS events were correlated with the prognosis of cancer including 1,324 positives related to cancer prognosis, and 1,597 negative correlated with cancer prognosis. 1172, 911, 837 AS events in 3UTR, 5UTR, and CDS region, respectively. Our ORF annotation analysis found 837 AS events in the CDS region would cause 417 in-frame and 420 frame-shift. (**Figure 4A**, **Table S4**).

**Figure 4.**
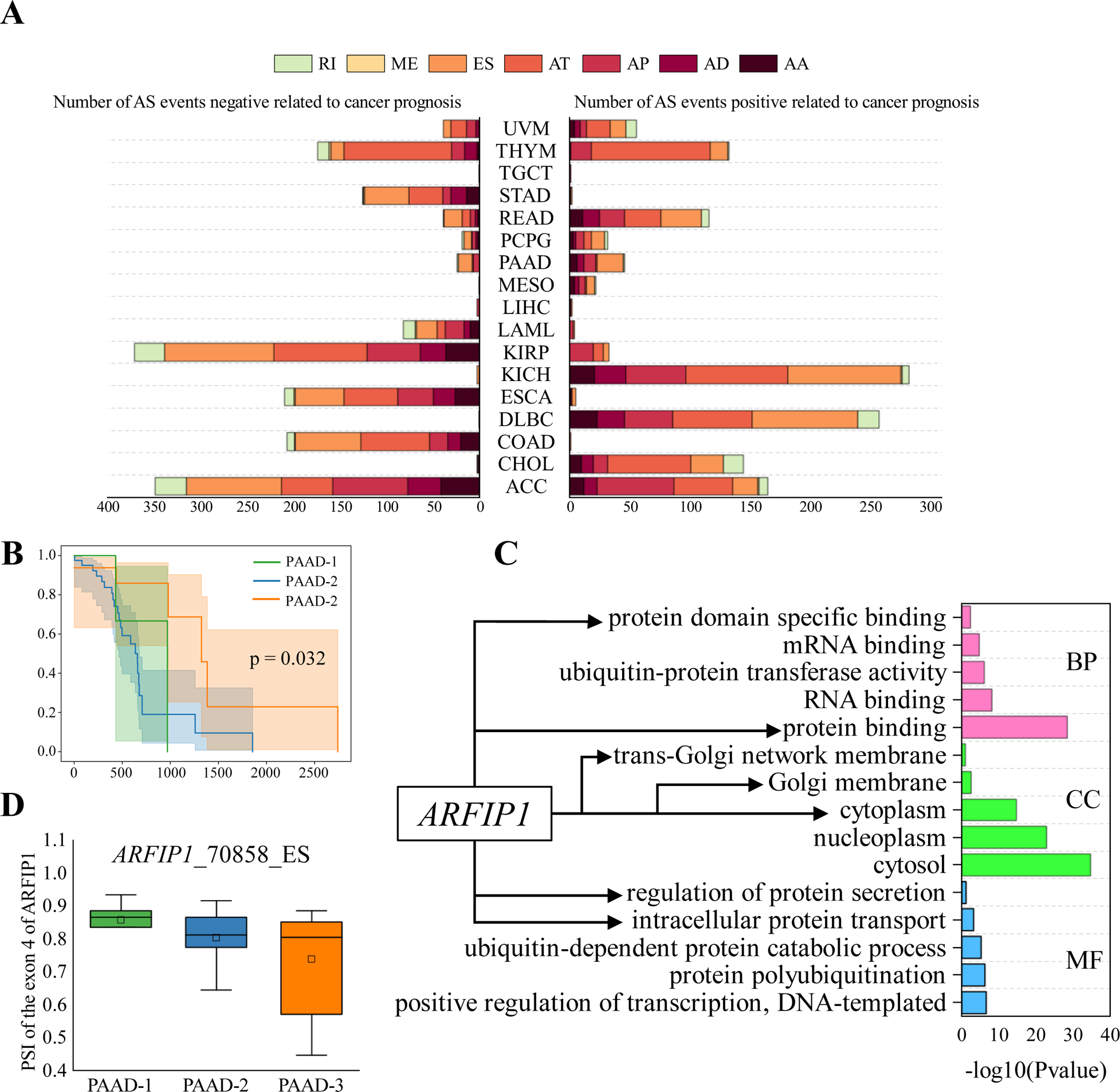
Results of the analysis of AS. **(A)** Pearson Analysis of AS events and Survival trend of subtypes, different colors represent different AS event types; (B) PAAD KM result; **(C)** GO enrichment PAAD subtype differential AS events; **(D)**Pearson analysis of AS event *ARFIP1*_70858_ES and subtype Survival trend.

An example is PAAD, The CLCluster identifies three subtypes of PAAD that show differential prognosis status (**Figure 4B**). 65 AS events were found to show different PSI values across PAAD subtypes and related to PAAD prognosis in 57 genes. The Gene Ontology enrichment analysis of the 57 genes showed that there were 158 significantly enriched terms, of which 28 were related to cellular components, 86 were related to biological processes, and 44 were related to molecular function. The Gene Ontology enrichment analysis showed that *ARFIP1* is enriched in the trans-Golgi network membrane (38), protein domain-specific binding (39), and other cancer-related GO terms **(Figure 4C)**. Of these events, the exon 4 (chr4: 153791904-153792000) of *ARFIP1* was negatively related to PAAD patients’ prognosis and showed differential PSI values across three subtypes **(Figure 4D)**. It has been reported that *ARFIP1* is ubiquitously expressed in human cancer cell lines (40). Our ORF annotation found the exon 4 skipping would cause the in-frame of the *ARFIP1* transcript which might affect the protein function and affected the PAAD progression.

Overall, these findings demonstrate the valuable insights that can be obtained by incorporating alternative splicing data into cancer subtyping analyses. By uncovering AS events that are associated with cancer subtypes and prognosis, researchers can gain a more comprehensive understanding of the molecular underpinnings of cancer and potentially identify novel therapeutic targets or biomarkers.

### 3.5 Screening for potential upstream RNA binds the protein of subtype-associated AS events

RNA-binding proteins (RBPs) play a crucial role in binding and regulating pre-mRNA splicing. In the context of aberrant splicing, there have been efforts to develop small molecules specifically designed to block RBP-mediated regulation, aiming to provide potential treatments for abnormal splicing patterns. To explore the regulation between RBP and AS event, we first performed ANOVA analysis and identified 291 RBP that showed differences between subtypes for 30 cancers. Furthermore, we performed correlation analyses between these RBPs expression and cancer prognosis stages identified from CLCluster, 91 RBPs related to patient survival were screened out from 8 cancers. Furthermore, 1752 RBP-AS pairs were found including 680 positive and 1071 negative RBP-AS regulations ((**Figure 5A, Table S5**). In these cases, RBPs may regulate associated alternative splicing events, and both the RBPs and alternative splicing events are linked to patient prognosis. Thus, these events can offer insights for the development of small-molecule drugs aimed at enhancing patient prognosis.

**Figure 5.**
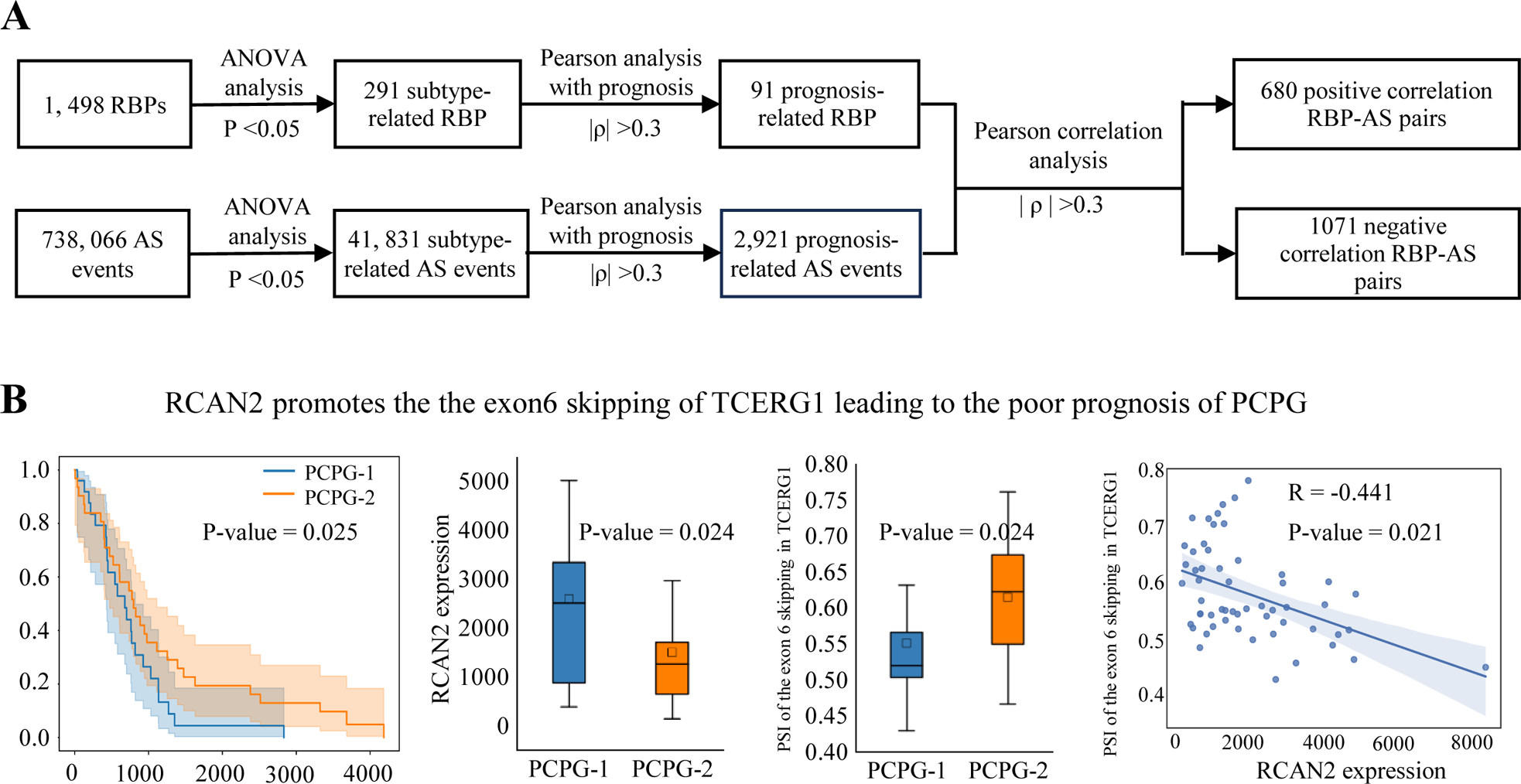
RBPs regulate the alternative splicing to affect patients’ prognosis. **(A)** The pipeline to identify the RBP regulates alternative splicing events in PCPG; **(B)** RCAN2 promotes the exon6 skipping of TCERG1 leading to the poor prognosis of PCPG. From left to right the figure shows KM result of PCPG; Gene expression of *RCAN2* in different subtypes of PCGA; PSI of the exon 6 of *TCERG1* in different subtypes of PCGA; Results of Pearson analysis of *RCAN2* and the exon 6 of TCERG1 in PCGA

As an example, the decreased expression of RNA-binding protein RCAN2 in PCPG has been associated with a favorable prognosis for patients. *RCAN2* can inhibit calcineurin activity by binding the catalytic subunit of calcineurin and leading to tumor proliferation(41). Furthermore, the association study found, that the increased expression of *RCAN2* leads to the skipping of exon 6(chr 5: 145847903 - 145847966) in *TCERG1* (**Figure 5B**). Conversely, inhibiting the expression of *RCAN2* can reverse the abnormal exon 6 skipping of *TCERG1*, potentially improving patient prognosis. These findings suggest that targeting the interaction between *RCAN2* and *TCERG1* could be a potential therapeutic approach to enhance patient outcomes.

## 4 Discussion and conclusion

Recent studies have found a significant correlation between Alternative splicing and cancer. The latest sequencing technologies have made a wealth of omic data accessible. As a result, there is now a much-awaited challenge to develop new techniques for integrating and analyzing multi-omics datasets, which can provide a better understanding of the intricate mechanisms of cancer.

In this work, we introduced alternative splicing data into cancer subtype clustering, which is the first time that alternative splicing data have been used in conjunction with other omics data for cancer clustering, and developed a comprehensive clustering method called CLCluster. This method uses redundancy-reducing contrast learning to achieve dimensionality reduction on multi-omics data and uses the mean-shift algorithm to obtain clusters automatically. Compared to other contrast learning algorithms that rely on negative examples, redundancy-reduced contrast learning does not use negative examples. This solves the problem that a contrast learning model will get trivial constant solutions and effectively prevents the model from collapsing. We utilized CLCluster to cluster 33 different types of cancers in the TCGA database. Our results showed that CLCluster performs better than the other seven advanced methods on most of the cancer datasets. Furthermore, we demonstrated that including AS data can enhance the clustering effect of cancer subtypes.

The identified splicing factors and biomarkers hold significant potential for various clinical applications. After obtaining the cancer subtypes from the CLCluster, we conducted further analysis. We screened 2,921 subtype-related exon skipping events that potentially caused the loss of function at the protein level and showed significant related to cancer prognostic progression by open reading frame annotation, protein function loss annotation, and RNA binding protein-associated alternative splicing regulation annotation and analysis. We can design antisense oligonucleotides to target these AS events’ splicing sites to avoid splicing abnormalities in cancer and by inhibiting the expression of these aberrantly expressed splicing factors, RNA can restore proper splicing and potentially impede tumor growth. In addition, we screened 1752 RBP-AS pairs, and the RBP in these pairs has a potential regulatory effect on AS events. We can use small molecule inhibitors for these pairs to disrupt the interaction between RBP and their related AS events, leading to altered splicing patterns and potentially inhibiting cancer cell growth.

While CLCluster introduces redundancy-reducing contrast learning into feature characterization, some limitations affect its performance. The downscaling and clustering functions in our model are performed separately. However, this may cause the model to overlook the linkages between the multi-omics data during clustering, which can reduce the effectiveness of the clustering to some extent. AS-based cancer therapies are already available. Our study screened for potential AS biomarkers, but whether these potential biomarkers are associated with AS events in existing therapies has not been validated, while the integration of information from relevant drugs with AS biomarkers has not been proposed. In future work, we will consider implementing an end-to-end model based on redundancy-reduced contrastive learning to accomplish both the downscaling and clustering tasks and adding drug information and treatment information to the network integration to build interaction networks to investigate the potential relationship between AS and drug treatment.

## Availability of data and material

The source codes are available at https://github.com/HIAHIA-catbus/CLCluster.

---

**Figure S1.**
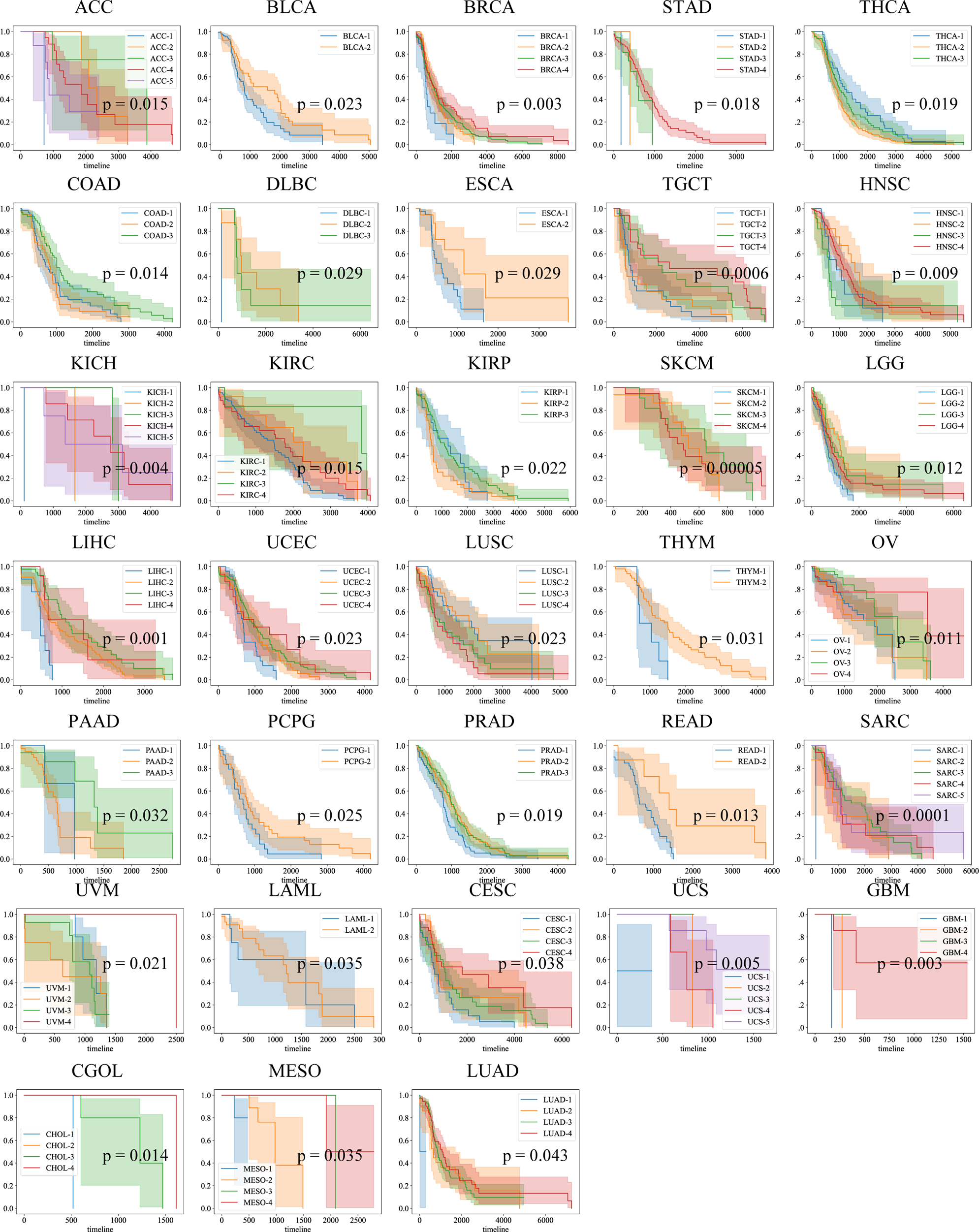
Survival analysis curves of CLCluster on 33 datasets. The different colors represent the grouping of samples according to the cluster labels output by CLCluster.

**Figure S2.**
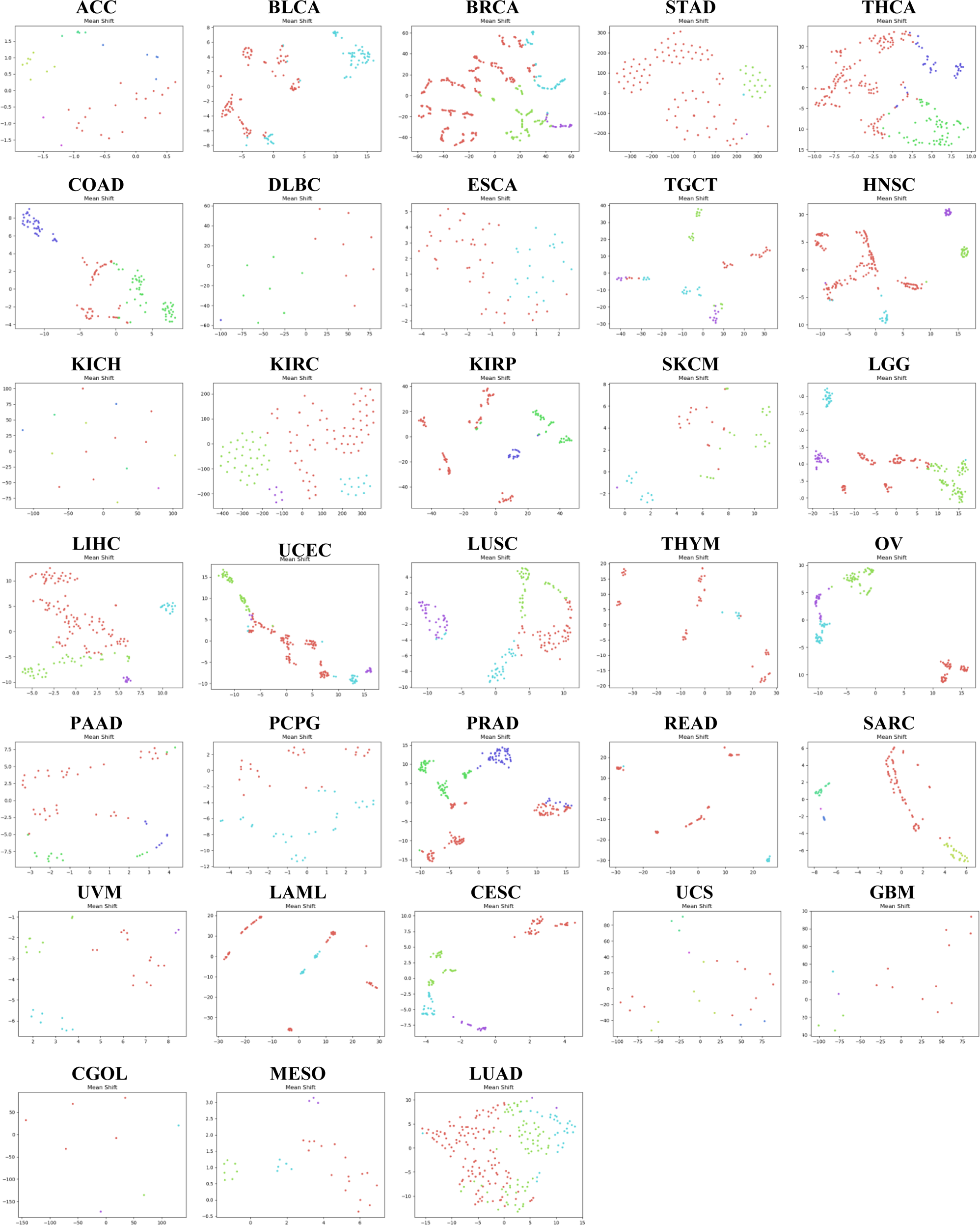
t-SNE of CLCluster on 33 datasets. The different colors represent the grouping of samples according to the cluster labels output by CLCluster.

